# Forward masking in the Inferior Colliculus: Dynamics of Discharge-rate Recovery after Narrowband Noise Maskers

**DOI:** 10.1101/2025.03.09.642170

**Authors:** Swapna Agarwalla, Afagh Farhadi, Laurel H. Carney

## Abstract

In forward masking the detection threshold for a target sound (probe) is elevated due to the presence of a preceding sound (masker). Although many factors are known to influence the probe response following a masker, the current work focused on the temporal separation (delay) between the masker and probe and the inter-trial interval (ITI). Human probe thresholds recover from forward masking within 150–300 ms, similar to neural threshold recovery in the IC within 300 ms after tone maskers. Our study focused on recovery of discharge rate of IC neurons in response to probe tones after narrowband gaussian noise (GN) forward maskers, with varying time delays. Additionally, we examined how prior masker trials influenced IC rates by varying ITI. Our findings showed that previous masker trials impacted probe-evoked discharge rates, with full recovery requiring ITIs over 1.5 s after 70 dB SPL narrowband GN maskers. Neural thresholds in the IC for probes preceded by noise maskers were in the range observed in psychoacoustical studies. Two proposed mechanisms for forward masking, persistence and efferent gain control, were tested using rate analyses or computational modeling. A physiological model with efferent feedback gain control had responses consistent with trends in the physiological recordings.

## I. INTRODUCTION

The perception of sensory events is influenced by spatial and temporal context (Fraisse, 1984; Hirsh, 1959; Lisker & Abramson, 1964; Stilp, 2020; Von & Pöppel, 1991). Forward masking is a phenomenon in which a preceding sound, the masker, causes a temporary reduction in the detectability of a subsequent target sound (Elliott, 1969; Plomp, 1964). Forward masking is hypothesized to have significant implications for perception of complex auditory stimuli, such as the suppression of echoes (Kaltenbach et al., 1993), the segregation of auditory streams (Fishman et al., 2004), and the enhancement of sensitivity to temporally structured stimuli, such as speech (Brosch & Schreiner, 1997).

Forward-masking experiments are common in psychoacoustical research that explores auditory thresholds, frequency discrimination, and the impact of context on sound perception. In forward masking experiments, researchers manipulate factors such as the duration, sound level, and frequency of both masker and target sounds (Jesteadt et al., 1982; Weber & Moore, 1981). Various peripheral and central mechanisms, including the middle-ear muscle reflex, neural adaptation, efferent innervation from the medial olivocochlear (MOC) bundle, and dynamic range adaptation, may collectively contribute to the phenomenon of forward masking (Jennings, 2021).

The concept of forward masking by noise is arguably more relevant to real-world situations than masking by pure tones. Noise stimuli may include the broad spectrum of frequencies found in many real-world sounds, such as the background noise in a crowded room or the rustling of leaves in the wind. Psychoacoustical studies have shown that probe-detection thresholds differ for tone and noise forward maskers (Weber & Moore, 1981). Physiological studies have traditionally focused on tone forward maskers at various places along the auditory pathway (auditory nerve: Relkin & Turner, 1988; dorsal cochlear nucleus: Kaltenbach et al., 1993; ventral cochlear nucleus: Shore, 1995; Ingham et al., 2016; medial nucleus of the trapezoid body: Gao & Berrebi, 2016). Recovery of the probe threshold in the inferior colliculus (IC) (Nelson et al., 2009) and auditory cortex (Alves-Pinto et al., 2010) is in a range (>40-50 dB) comparable to that observed psychoacoustically. Recovery of physiological thresholds remains unexplored for noise maskers. The primary goals of the current work were to quantify the recovery from masking by narrowband gaussian-noise (GN) maskers in the IC by varying the delay between the masker and probe and to investigate the impact of the inter-trial interval (ITI) on responses to probe tones in subsequent trials. This approach allowed for estimating carryover effects of maskers, across trials, particularly for shorter ITIs. The effect of ITI on physiological forward-masked thresholds was also estimated.

The results were used to test two hypotheses for mechanisms underlying forward masking. Analyses of the rates were used to test the long-standing persistence hypothesis (Moore et al., 1988; Oxenham, 2001). Additionally, previous modeling studies have supported the hypothesis that forward masking may be partially explained by cochlear gain control via the medial olivocochlear system (Brennan et al., 2023; Maxwell et al., 2024). Motivated by that work, responses of physiological models with and without efferent gain control (Farhadi et al., 2023) were compared to physiological recordings.

## II. MATERIALS AND METHODS

### A. Animals and Neural Recordings

The study was approved by the University of Rochester Committee on Animal Resources and was carried out in adherence to the National Institutes of Health guidelines. Dutch-Belted female rabbits *(Oryctolagus cuniculus),* aged 1-5 years were studied. Rabbits had normal hearing, verified using distortion-product otoacoustic emissions (DPOAEs) (Whitehead et al., 1992) and IC neural thresholds. Surgical procedures and neural-recording techniques have been reported previously (Fan et al., 2021, 2022; Mitchell et al., 2023). Briefly, in anesthetized rabbits, aseptic surgeries were performed to establish chronic access to the midbrain. Either intramuscular ketamine (66 mg/kg) and xylazine (2 mg/kg) or ketamine (66 mg/kg) and dexmedetomidine (0.10-0.15 mg/kg) anesthesia and subcutaneous meloxicam (0.2 mg/kg) analgesia were used for surgical placement of a custom 3D-printed plastic headbar, used to immobilize the head during recording sessions and to hold a microdrive. After recovery, during a second surgery, a 2-mm-diameter craniotomy was made and a manual microdrive (Neuralynx 5-drive, Boseman, MT) was positioned. The microdrive was used to advance four chronically implanted tetrodes made with 18-µm platinum-iridium wire and plated with platinum-black to achieve impedances of 0.1-0.5 MΩ.

Extracellular single-unit action potentials were recorded, using a 30-kHz sampling rate, in the IC of awake rabbits in daily 2-hour recording sessions in a sound-attenuated booth (Acoustic Systems, Austin, TX). Tetrodes were advanced slowly through the IC over the course of several weeks, after which microdrive-replacement surgery was performed to change tetrodes and recording sites. For neural recordings, a headstage-amplifier recording system and software (RHD2000, Intan Technologies, LLC., Los Angeles, CA) were used, including an analog high-pass filter (1st-order, 150 Hz), an analog, Butterworth, low-pass filter (3rd-order, 7.5 kHz), and a digital high-pass filter (1st-order, 300 Hz), followed by 16-bit analog-to-digital conversion.

The action-potential waveforms and times were extracted from neural recordings using custom MATLAB code (Mathworks, Natick, MA). To isolate recordings from single neurons, the voltage recordings were digitally bandpass filtered from 300 - 3000 Hz, and a voltage threshold was typically set at four standard deviations above the mean of the voltage signal, adjusted, if necessary, after visual inspection. Action potentials were categorized into clusters, assumed to represent distinct neurons, based on the slope of the waveform repolarization (Schwarz et al., 2012). Recordings were assumed to be from isolated neurons when fewer than 2% of the interspike intervals were less than 1 ms, and when multiple clusters were well separated, based on the cluster separation metric (Schwarz et al., 2012).

### B. Stimuli

Stimuli were created using custom MATLAB code and delivered by an audio interface (16A, Mark of the Unicorn, Cambridge, MA) at a sampling rate of 48 kHz with a 16-bit digital-to-analog converter (DAC3 HGC, Benchmark Media Systems, Inc., Syracuse, NY). Sound stimuli were presented by earphones (ER2, Etymotic Research, Inc., Elk Grove Village, IL, or BeyerDynamic, Heilbronn, Germany) connected to custom-made ear molds (Dreve Otoform Ak, Unna, Germany).

Before each session, the frequency response (magnitude and phase) of the acoustic system was calibrated using ER-7C or ER-10B+ probe-tube microphones (Etymotic Research, Inc., Elk Grove Village, IL) to specify a pre-emphasis filter that was applied to all stimuli. The frequency responses were tapered outside the 0.1-18-kHz frequency range, using a cubic-spline interpolation (MATLAB function *csape*) to create a smooth curve through a set of data points with piecewise cubic polynomials, with zero slope at the onset and offset of each stimulus, to minimize the ringing in the silent period between masker and probe stimuli.

Recording sessions began with the presentation of several stimuli to characterize the recording sites. Response maps, or measurements of average rate vs. frequency at several sound levels, were used to identify the characteristic frequencies (CFs) at which neurons were most sensitive. Response maps were based on average responses, including onset responses, to three repetitions of a randomized sequence of 200-ms diotic tones, spanning frequencies from 250 Hz to 16 kHz, at levels of 10, 30, 50, and 70 dB sound pressure level (SPL), with 5 steps per octave. Response maps were labeled based on visual inspection as follows (Ramachandran et al. 1999): Type V, an excitatory area that increased in bandwidth with sound level; Type I, a narrowband region of excitation flanked by inhibition at lower and/or higher frequencies; Type O, excitation at low stimulus levels bounded by inhibition at higher levels. Onset and Offset Types were dominated by onset and offset responses to the tones. Unusual Types had response maps that did not fit into any of the above categories.

Modulation transfer functions (MTFs) were also measured, consisting of average response rates (excluding the first 25-ms) to 100% sinusoidally amplitude-modulated (AM) wideband noise spanning 100 Hz–10 kHz, with 1-s duration, including 50-ms cos^2^ on/off ramps, across a modulation-frequency range of 2-600 Hz, with 3 steps per octave. AM stimuli were presented diotically for 5 repetitions at a spectrum level of 30 dB SPL (overall level of 70 dB SPL). MTFs were classified based on the average rates across a range of modulation frequencies compared to the rate in response to an unmodulated noise. MTFs were labeled band-suppressed (BS) or band-enhanced (BE) when rates were significantly reduced or increased, respectively, compared to the unmodulated response rate (unpaired t-test, p<0.05) for at least two successive modulation frequencies. MTFs for which rates displayed both enhancement and suppression within different modulation-frequency bands were labeled hybrid, with two subcategories: BE-BS were enhanced at lower modulation frequencies and suppressed at higher frequencies; BS-BE neurons were suppressed for lower modulation frequencies and enhanced at higher frequencies. Any neuron failing to exhibit significant differences in comparison to unmodulated rates (unpaired t-test, p<0.05) across at least two consecutive modulation frequencies was categorized as flat.

#### 1. Experiment 1: Forward-masking paradigm with varied time delay between masker and probe

The goal of this experiment was to quantify the recovery of the average-rate response to a probe tone preceded by a narrowband-noise masker. Stimuli were similar to those used in the psychoacoustic study of Brennan et al. (2023), except that here the masker level was set to 70 dB SPL, and the probe level was fixed at 60 dB SPL, which would be above detection threshold for listeners with normal hearing. GN maskers had a center frequency matched to the CF of a neuron, a bandwidth equal to 1/3 of the equivalent rectangular bandwidth of the auditory filter (ERBn, Glasberg & Moore 1990) at the center frequency, and a duration of 400 ms, including 5-ms cos^2^ on/off ramps. Masker waveforms varied across trials. Control conditions consisted of trials in which only the probe was present (i.e., the masker was absent). The masker was followed by a probe tone at a frequency matched to CF, 10-ms duration, including 5-ms cos^2^ on/off ramps. The zero-amplitude delay between masker offset to probe onset was 0, 0.025, 0.075, 0.150, 0.250, 0.5, 1.0, 1.5, or 2 s.

The stimuli were presented diotically in random order with an ITI of 2 s. Each stimulus condition was presented 20 times.

#### 2. Experiment 2: Forward-masking paradigm with varied ITI between trials

The goal of this experiment was to test for a potential prolonged impact of maskers on subsequent trials by measuring responses to unmasked probe tones at various ITIs following trials with maskers.

The ITI was defined as the interval between the offset of the probe and the onset of the masker in the following trial. For trials without a masker, a silent interval was included that matched the combined duration of the masker and the gap between the masker and probe, ensuring that all trials for a given delay condition had the same overall duration. The stimuli for this experiment were the same as in Experiment 1, except that i) the delay between the GN masker and probe was fixed at 0.25 s, ii) the ITI of 0.5, 1 or 1.5 s were presented in blocks, iii) For carryover effect from previous trials, responses to unmasked probes on trials that were preceded by masked trials were compared to responses to unmasked probes in the control block. The control block consisted of only unmasked probes. Three probe levels (60, 65, and 70 dB SPL) were used to quantify the carryover effect. iv) To estimate neural thresholds for probe detection, for conditions with or without a masker, no-probe responses were compared to responses for which probe level was randomly varied from 5-70 dB SPL in steps of 5 or 10 dB. Different trial types were randomly interleaved during recording sessions. For each ITI condition, there were 20 no-probe trials and 20 trials at each probe level. Each block included different probe levels, but the overall stimulus duration and inter-trial interval (ITI) were fixed within a block. For example, if the stimulus consisted of a masker (0.400 s) + delay (0.025 s) + probe (0.010 s) + ITI, the total stimulus duration would be 0.435s + ITI. In conditions without a masker, the same duration was maintained by filling the masker period with silence, followed by the ITI of either 0.5, 1, or 1.5 s. Spontaneous activity was measured from a 0.4-s window immediately before the start of the subsequent trial (Nelson et al., 2009).

### C. Analysis of physiological recordings

For Experiments 1 and 2, neurons that had a significant change in average rate in response to the unmasked probe compared to spontaneous activity (t-test, p < 0.05) were considered for further analysis. The discharge rate in response to the probe tone was computed over a 30-ms window starting from the onset of the probe tone. A response was considered to be recovered if there was no significant difference between the masked and unmasked probe rates (t-test, p > 0.05). Response recovery from masking was assessed by calculating the ratio between the average discharge rates in response to masked and unmasked probe tones.

For Experiment 2, single-neuron detection thresholds for the probe were estimated using receiver operating characteristic (ROC) analysis (Egan, 1975) with a criterion of 70.7% correct. The response to no-probe and probe trials were analyzed using the same time window following the masker, 30- ms window starting from the onset of the probe tone. Additionally, a population threshold was estimated using a maximum likelihood-based pattern-decoder analysis (Day and Delgutte, 2013; Jazayeri and Movshon, 2006). The detection threshold for the population was evaluated using single-trial responses that were optimally weighted assuming independent, Poisson-distributed spike counts for each neuron and stimulus. The decoder computed the likelihood of each stimulus alternative (with and without probe, at different sound levels) by generating 1000 population responses for masked (or unmasked) conditions. For each draw from the population of neurons, the logarithm of the likelihood of each alternative was calculated as

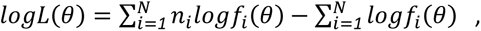

where *n*_*i*_ is the randomly drawn spike count, *f*_*i*_(*θ*) is the mean spike count for stimulus *θ* of the *i^th^* neuron, and *N* is the total number of neurons. The first term is an optimally weighted sum of spike counts and the second term is the sum of mean spike count across the population. Detection performance was measured for each stimulus alternative (with and without probe) as the proportion of population draws for which the likelihood of the correct stimulus condition surpassed that of the alternative. The population threshold was defined as the minimum sound level at which the decoder achieved classification performance exceeding 70.7% correct.

### D. Phenomenological Model

#### 1. Estimating MOC efferent model parameters for cat tuning

Simulations were made using computational models for auditory-nerve (AN) responses, either without (Zilany et al., 2014) or with (modified version of Farhadi et al., 2023) efferent control of cochlear gain. The Zilany et al. (2014) model is based on peripheral tuning in cat. The Farhadi et al. (2023) subcortical model includes peripheral tuning in human; here the model was modified for cat tuning, based on Zilany et al. (2014), and efferent parameters were estimated using rabbit data.

Sustained stimulus fluctuations may progressively reduce cochlear gain via medial olivocochlear (MOC) efferent feedback, leading to a delayed (by 100–200 ms) increase in Band-Enhanced (BE) IC neuron firing rates in response to AM noise. This rate increase occurs because reduced cochlear gain enhances the modulation depth of inner-hair-cell responses, which ultimately increases IC BE responses until efferent control stabilizes. Notably, IC BE responses in awake rabbits to wideband sinusoidal-AM gaussian noise in awake rabbits (Carney et al., 2014) have rates that increase over time, consistent with MOC dynamics (Farhadi et al., 2023).

The efferent model parameters (time constant of the MOC pathway, *τ*_MOC_; rational-function parameter of the MOC input-output function, *B*; and the scaler on the IC input to the MOC, *K_ic_*) were estimated by fitting the AN model to a set of rabbit IC BE responses to wideband sinusoidally AM Gaussian noise (from Carney et al., 2014), using procedures described in Farhadi et al. (2023) (Fig. 1). The model with efferent gain control (dot-dashed red) based on the mean values of the distributions of model parameters (*τ*_MOC_ = 193.8 ms, *B* = 0.02, and *K_ic_* = 44.7) provided a better prediction of the neural response (solid black) over the time course of the AM wideband Gaussian noise (Fig. 2) than the model without efferent gain control (dashed blue). The IC BE rate increased over the steady-state part of the response (gray time window, excluding onset) for most neurons with IC BE MTFs (Farhadi et al., 2023).

**Figure 1:**
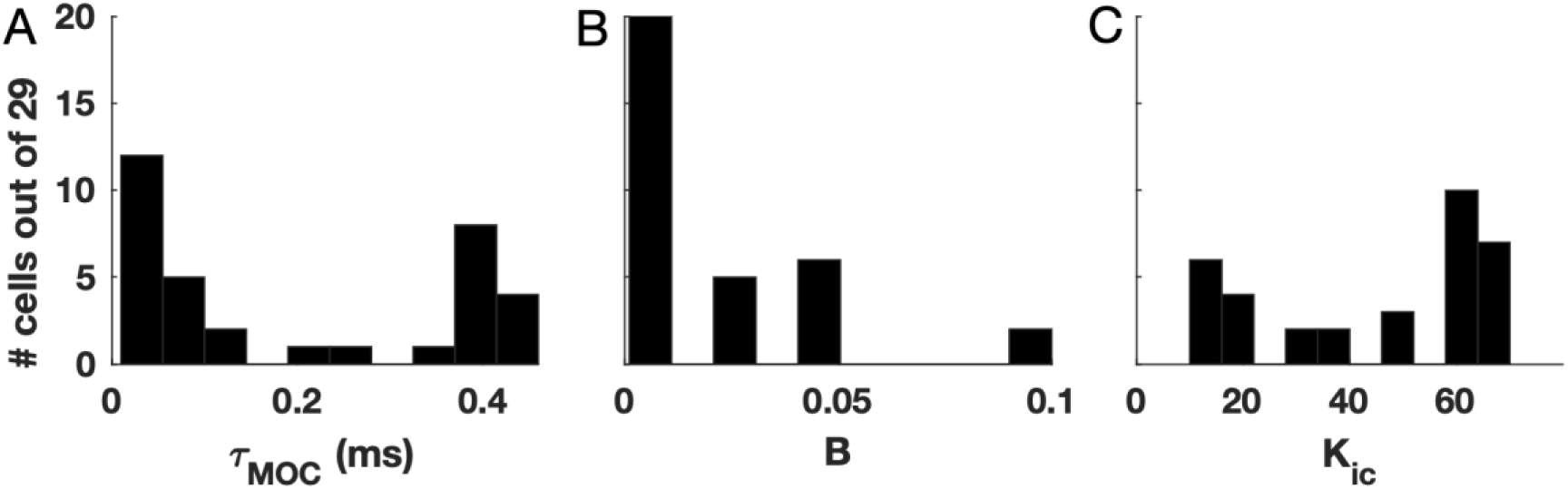
Distributions of the efferent pathway parameters for the subcortical model with cat peripheral tuning, fitted to responses to wideband AM noise in rabbit IC. A) Time constant, *τ*_MOC_, B) Rational-function parameter of the MOC input/output function, *B*, and C) Scaler applied to the IC input to the MOC, *K_ic_*. Parameter fits were based on the means of these distributions for 35 IC BE neurons (from Carney et al., 2014) with increasing rates during the wideband AM noise stimuli and BMFs near the stimulus AM frequency.

**Figure 2:**
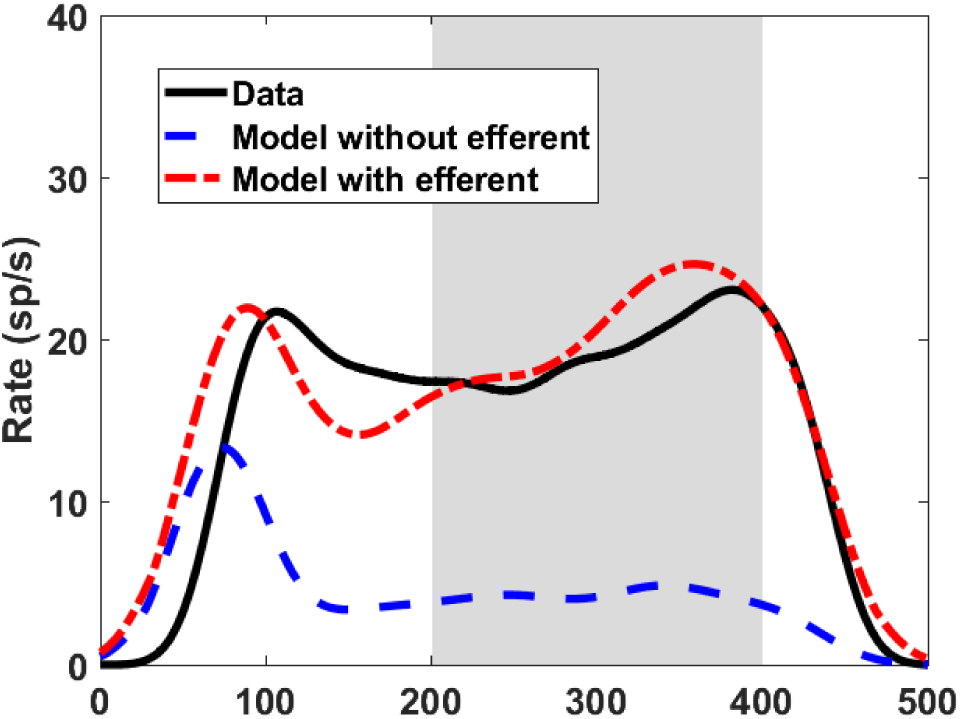
A) Example IC neuron response (CF = 3.5 kHz, BMF = 80 Hz) in awake rabbit (black) to wideband sinusoidal-AM noise with modulation depth of -10 dB. The CF and BMF of the subcortical model were matched to the example neuron. Median values for the model parameters (*K_ic_* = 55, *τ*_MOC_ = 85 ms, and *B* = 0.01) were used for the model with efferent feedback.

#### 2. Predicting physiological responses to narrow-band forward masking across varying masker-probe delays

The IHC-AN synapse model was updated to use a fast power-law-adaptation implementation (Guest & Carney, 2024). Simulations used a sampling frequency of 100 kHz. The AN models provided the input to the same-frequency inhibition-excitation (SFIE, Carney & McDonough, 2019; Nelson & Carney, 2004) model of BE IC neurons, with best modulation frequency (BMF) set to 64 Hz, the median BMF reported by Kim et al. (2020) for rabbit. Probe-tone detection thresholds were determined using instantaneous discharge rates from BE IC model neurons. Spontaneous activity of the model IC neurons resulted from AN spontaneous activity, driven by fractional gaussian noise (Zilany et al., 2014). The stimuli used for simulation and the analysis procedures were the same as those in Experiment 1. The code for the model and data analysis will be made available at https://osf.io/n4m5b.

## III. RESULTS

### A. Experiment 1

#### 1. Recovery of discharge rate as a function of delay between GN masker and probe

Recordings were made from 80 well-isolated units in 2 animals. A schematic of the stimuli and representative responses are shown in Fig. 3A-C. The recovery from masking for each neuron was quantified by the ratio of discharge rates in response to masked vs. unmasked probes, referred to as the recovery ratio. As observed in previous psychophysical (Brennan et al. 2023; Jesteadt et al., 1982; Moore & Glasberg, 1983; Weber & Moore, 1981) and physiological (Alves-Pinto et al., 2010; Gao & Berrebi, 2016; Ingham et al., 2016; Kaltenbach et al., 1993; Nelson et al., 2009; Relkin & Doucet, 1991; Shore, 1995) studies of forward masking, recovery from masking increased with delay (Fig. 4A). Neurons that exhibited a recovery ratio greater than 1 at zero delay had strong offset responses to the masker. A neuron was considered fully recovered if the distributions of masked and unmasked probe discharge rates were not significantly different (t-test, p > 0.05).

**Figure 3:**
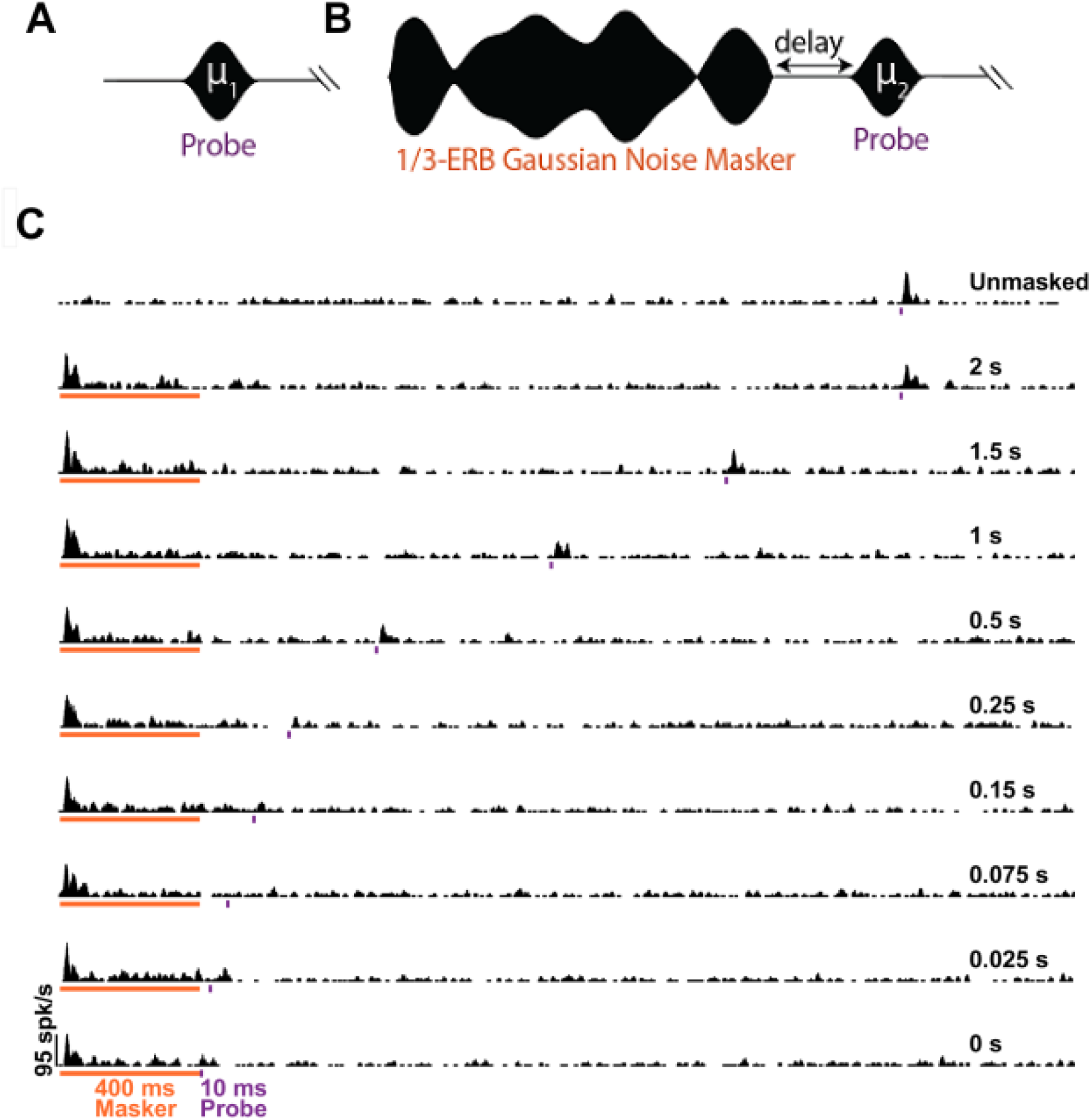
A, B) Schematic of the stimuli for Experiment 1. The control condition with an unmasked probe (left) and the test condition with a masked probe (right). The delay between GN masker and probe was varied. C) Representative peristimulus time histograms (PSTHs) in response to unmasked probe (top) and for GN maskers followed by probes at various delays. The orange horizontal line indicates the masker presentation time (400-ms duration); the violet markers indicate the timing of the 10-ms duration probe tones. Probe delays are indicated at the right of each PSTH.

**Figure 4:**
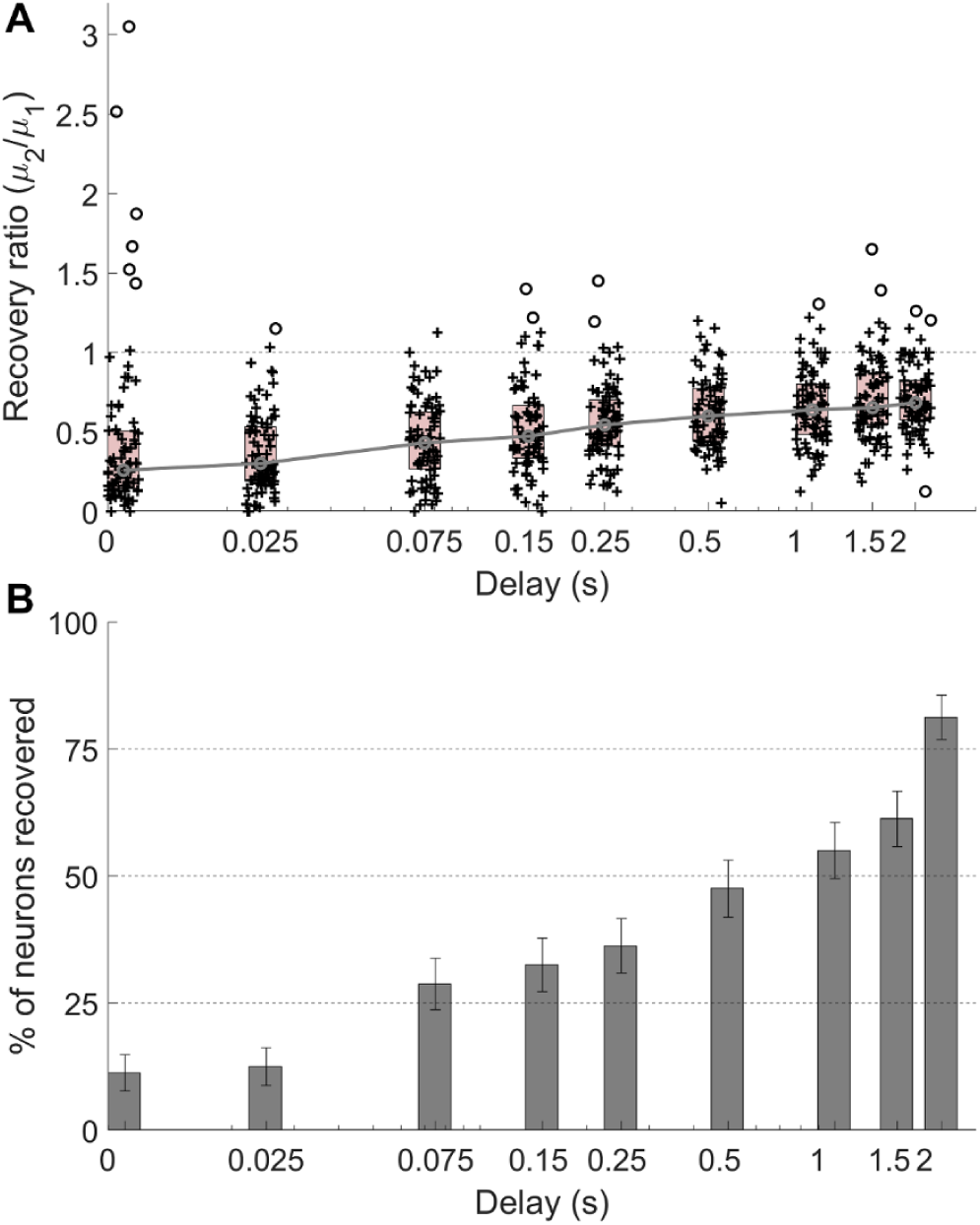
A) Recovery ratio as a function of delay. Each ‘+’ symbol indicates a neuron; outliers are represented by ‘o’ symbols. The shaded rectangle represents the interquartile range; the medians are joined by a gray line. The horizontal gray dashed line at 1 indicates full recovery from masking. B) The percentage of neurons (+/- standard error) that were fully recovered from masking at each delay.

Previous physiological studies with tone maskers show that a delay of 150-300 ms is sufficient for the recovery of neurons for masker levels 30-60 dB above neural thresholds estimated for the unmasked probe (medial nucleus of the trapezoid body, rat,Gao & Berrebi, 2016; IC, marmoset, Nelson et al., 2009; auditory nerve, chinchillas, Relkin & Doucet, 1991; ventral cochlear nucleus, guinea pig, Shore, 1995). In the current study, longer delays were required for recovery of probe response after GN maskers at 70 dB SPL: approximately 52% of neurons were not recovered from masking for a probe delay of 0.5 s, and almost 20% were still not recovered for a delay of 2 s (Fig. 4B). For most neurons, the 70 dB SPL masker level was 40 dB or less above the RM threshold at CF, thus, a relatively short recovery would be expected. Previous psychoacoustic studies suggest that masker duration also affects the recovery from masking (Kidd & Feth, 1982; Oxenham & Plack, 2000). The relatively long recovery after the GN forward maskers might be partly due to the relatively long, 400-ms-duration maskers used here, chosen to match those in a psychophysical study (Brennan et al., 2023), as compared to the shorter-duration tone maskers used in previous physiological studies, typically in the range of 40-200 ms (50 ms, 102 ms, Alves-Pinto et al. 2010 in auditory cortex of guinea pig; 200 ms, Gao & Berrebi, 2016 in medial nucleus of the trapezoid body of rat; 40 ms, Kaltenbach et al., 1993 in dorsal cochlear nucleus of hamster; 200 ms, Nelson et al., 2009; 102 ms, Relkin & Doucet, 1991 in auditory nerve of chinchillas; 50, 100 ms, Shore, 1995 in ventral cochlear nucleus of guinea pig).

A decrease in probe response does not necessarily indicate forward masking because a reduction in spontaneous rate following the masker could enhance the detectability of the probe, as was pointed out in studies of forward masking at the level of the auditory nerve (Relkin and Turner, 1988; Turner, Relkin, Doucet, 1994). To investigate this possibility, the discharge rate between the masker and the probe for different delays (excluding the 0-ms delay) was calculated and compared to the spontaneous rate measured 400-ms before stimulus onset of the next trial (Nelson et al., 2009). Our analysis revealed that only 10% of neurons exhibited suppression of rate with respect to spontaneous rate during the interval between the masker and probe for the 25-ms delay condition. The change in spontaneous rate gradually diminished to zero by a delay of 1.5-s. It is important to note that, in contrast to the auditory nerve, most IC neurons have very low baseline spontaneous rates (e.g., 96% of neurons have spontaneous rates less than 20 spikes/s).

#### 2. Examining Recovery from Masking: Spontaneous Activity, CF, Masker-Response Type

The correlations between recovery from masking and response features such as spontaneous activity and CF were more pronounced for shorter delays between the masker and probe (cochlear nucleus, Shore, 1995). Thus, responses to probes delayed by 0 ms were examined to investigate correlations between recovery ratio and spontaneous activity, CF, and masker responses. Pearson correlations (Fig. 5A) between spontaneous activity and recovery ratio at the 0-ms delay were not significant. Higher-CF neurons had significantly less recovery from masking at the 0-ms delay (Fig. 5B).

**Figure 5:**
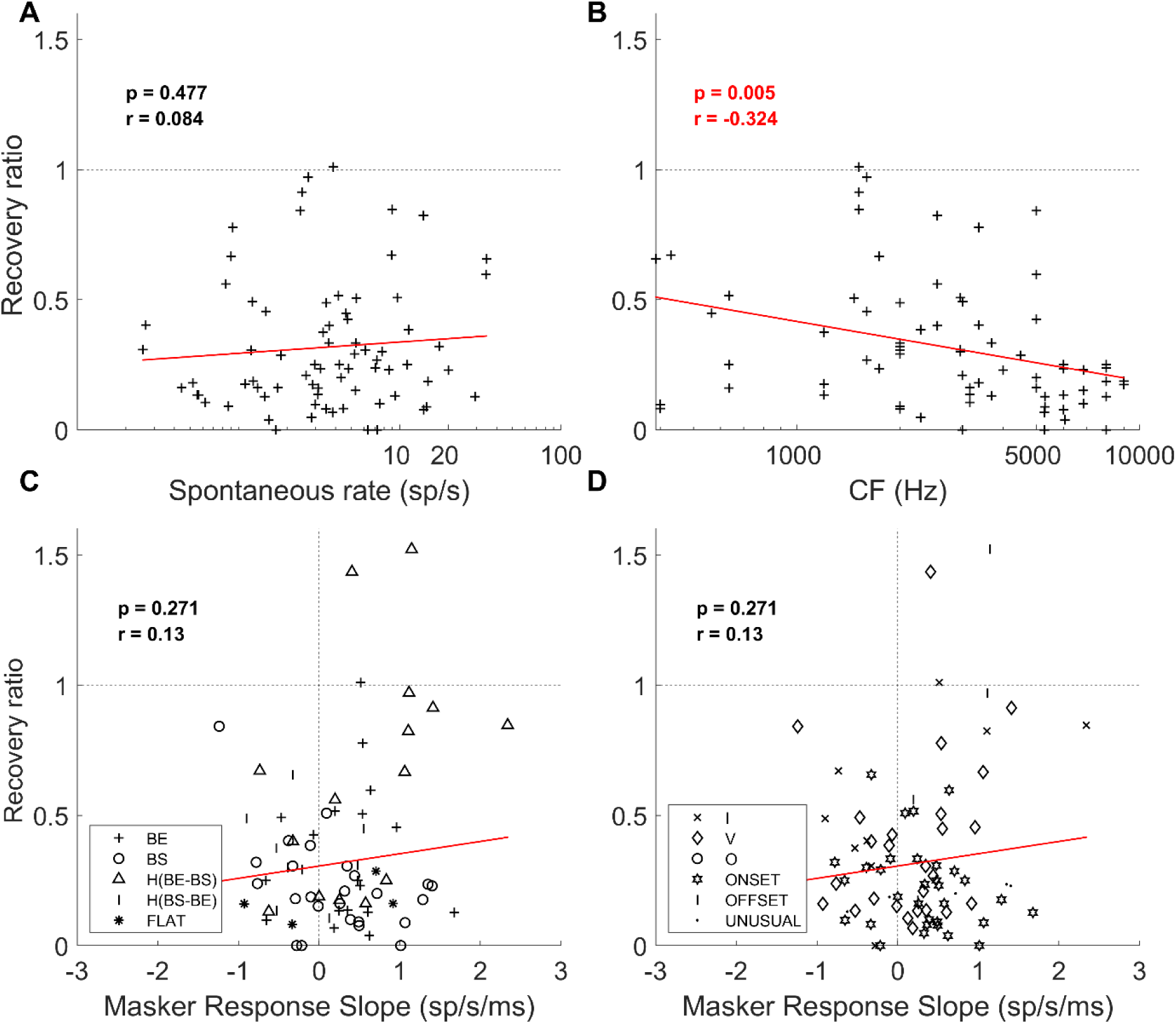
Recovery ratio plotted for responses to stimuli with 0-ms delay between GN masker and probe as a function of A) spontaneous rate, B) CF, C) Masker-response slope, based on a line fitted to the PSTH (binsize 1 ms) from 100 to 400 ms after masker onset. Different symbols represent different MTF types. D) Same data as in C, but different symbol shapes represent different RM types. The horizontal gray dashed line indicates full recovery from masking. For all subplots, red solid lines are linear fits to the data. Pearson correlation, *r*, and associated p-values are reported in each panel. Outliers (plotted in Fig. 4A) were not included in these plots or in the regressions.

To test the hypothesis that differences in the time course of the masker response affect the recovery of a neuron from masking, the masker responses were assessed by normalizing the PSTH by the peak bin (1-ms binsize) and calculating the slope of a line fitted to the normalized PSTH from 100 to 400 ms after masker onset. A positive slope indicated a buildup of the masker response over time, whereas a negative slope suggested adaptation of the masker response over time. There was not a significant correlation between the masker-response slope and recovery ratio (Fig. 5C, D).

Qualitatively, the recovery from masking did not appear to be related to MTF type (Fig. 5C) or RM type (Fig. 5D).

To ensure that the analyses of responses to probes delayed by 0 ms were not influenced by persistence of the masker response, similar analyses were done for a delay of 25 ms. The findings were similar to those at 0 ms, with no significant correlation between the recovery from masking and spontaneous activity (p > 0.05) or masker-response slope (p > 0.05), and a significant correlation between masker recovery and CF (p < 0.05).

### B. Experiment 2

#### 1. Effect of maskers on probe responses in subsequent trials without maskers

ITI was not a focus of previous physiological studies of forward masking, and the potential for persistent, or “carryover,” masking from previous trials has not been explored. The ITIs used in forward-masking studies with tone maskers differ across studies (auditory cortex, guinea pig: 1-1.6 s, Alves-Pinto et al., 2010; medial nucleus of trapezoid body, rat: 1 s, Gao & Berrebi, 2016; ventral cochlear nucleus, guinea pig: 450 ms, Ingham et al., 2016; IC, marmoset: ∼470-790 ms, Nelson et al. 2019; auditory nerve, chinchillas: 273 ms, Relkin & Doucet, 1991; ventral cochlear nucleus, guinea pig: 300 ms, Shore, 1995). We measured the average effect of masked trials on immediately following unmasked-probe trials in 96 units from 2 animals, although all ITIs was not studied in all units. Stimuli were presented with masker level fixed at 70 dB SPL and probe level at 60, 65, or 70 dB SPL; some units were studied at more than one level. Trials with masked and unmasked probes were presented in a pseudorandom manner, with 50% of trials of each type (Fig. 6A). The control block consisted of only unmasked probes (Fig. 6B). The ratio between the unmasked-probe rates in control blocks (**μ**_2_, Fig. 6B) and the unmasked-probe rates in trials that were preceded by masking trials (**μ**_1_, Fig. 6A), is shown for different ITIs (Fig. 6C). As ITI increased, the ratio trended towards 1, where a value of 1 would indicate no effect of the previous masker trial. A 1-way ANOVA followed by a post-hoc Tukey-Kramer multiple-comparison analysis showed that these ratios differed significantly for ITI values of 0.5 s and 1.5 s (p < 0.05), but not for other pairwise comparisons (p > 0.05). The percentage of neurons with ratios that were not significantly different from 1 is shown in Fig. 6D (t-test, p > 0.05). Approximately 65% of neurons were recovered from masking on previous trials for ITI values of 0.5 s and 1 s. For an ITI of 1.5 s, the percentage of recovered neurons increased to 78%. In other words, 22% of neurons had responses to the probe tone that were still reduced by maskers on previous trials for an ITI of 1.5 s. To quantify the effect of carryover on the masker, the onset response (50-ms window from masker onset) and the average rate during the masker were compared across ITIs. No significant differences were observed (1-way ANOVA, p > 0.05).

**Figure 6:**
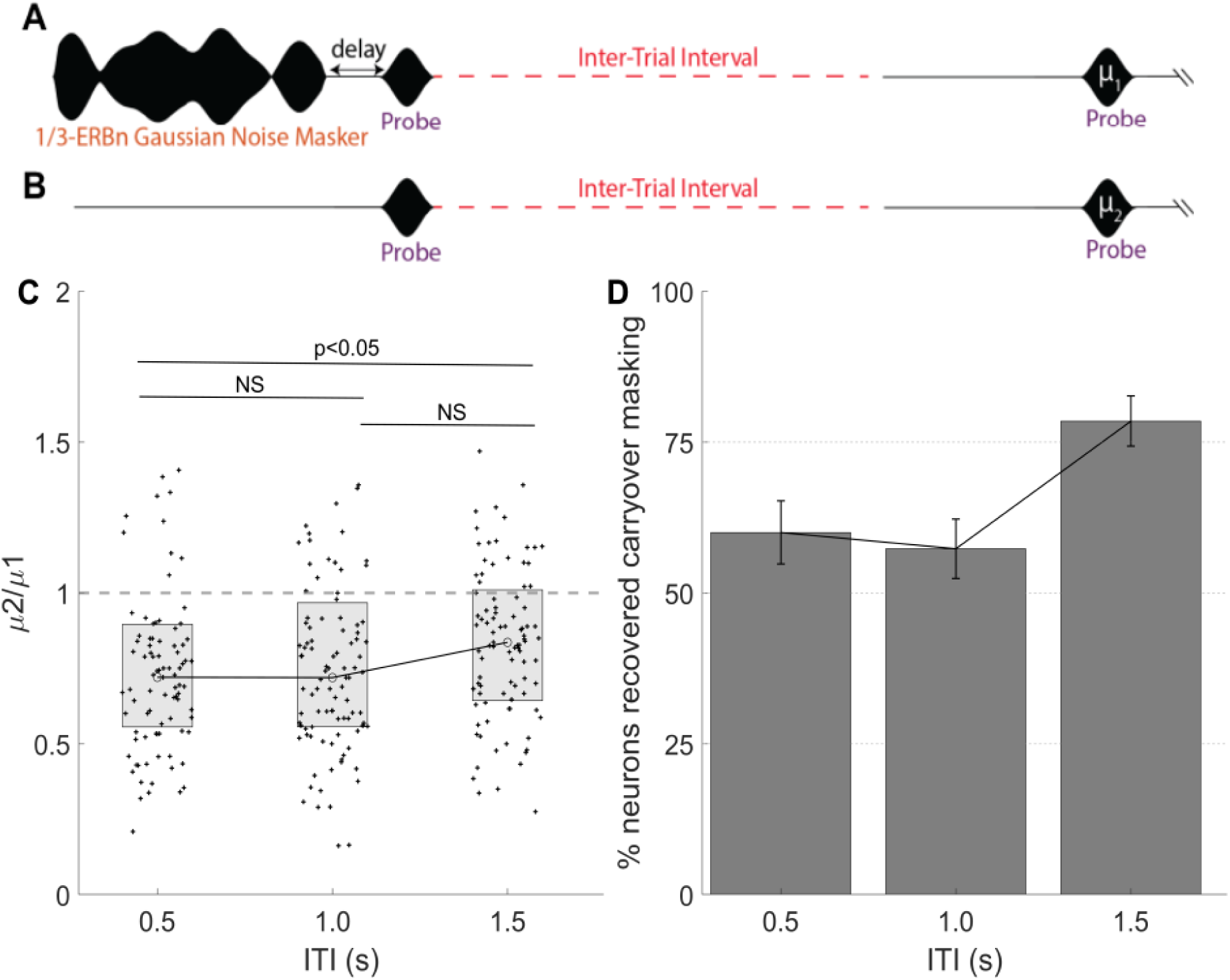
A & B) Stimuli for Experiment 2. A) Trials with and without maskers were presented in a pseudorandom manner. Unmasked trials that were immediately preceded by masked trials were included in this analysis. B) The control block comprised only unmasked probes. The ITI (red) between the trials varied across blocks. C) Ratio of discharge rates for unmasked probes in control block to those following masking trials as a function of ITI (n=87, 96 and 93 for ITIs 0.5, 1.0, 1.5, respectively). D) Percentage of neurons (+/- standard error) with ratios that were not significantly different from 1 (i.e., unmasked probe responses were not different between condition A and B), as a function of ITI.

#### 2. Masked and Unmasked Probe Thresholds

Neural thresholds could be estimated for both unmasked and masked probes (Fig. 7A) for 19, 20 and 21 units in 2 animals, for ITIs of 0.5, 1, and 1.5 s, respectively; ITI was varied in blocks. In an additional 31 units, the masked threshold exceeded the range of levels tested (maximum SPLs tested were typically 70- or 60-dB SPL, except for three units with a maximum of 50 dB SPL); these units, despite being strongly masked, were not included in this analysis. PSTHs for masked and unmasked probe responses of one neuron are shown in Fig. 7B; black arrows indicate the lowest probe levels at which responses were significantly above the spontaneous rate for either masked or unmasked probes. The average discharge rate for the same neuron is shown in Fig. 7C for unmasked (solid lines) and masked (dashed) probes. The rate elicited by the probe was compared to that measured over an equivalent temporal window when no probe was presented. To estimate threshold, responses to pairs of probe and no-probe trials drawn from all the repetitions were compared, and correct detection was determined when the probe response was greater than the no-probe response. For cases that did not differ in probe rate, correct detection was determined 50% of the time, based on a random draw. An ROC analysis (Egan, 1975) was used, with criteria ranging from 0 to maximum rate; the area under the curve was calculated for each probe level to estimate percent correct. This process was repeated for all probe levels to construct neurometric functions (Fig. 7D). Neural detection thresholds were estimated by finding the minimum probe level that exceeded 70.7% correct (Fig. 7D).

**Figure 7:**
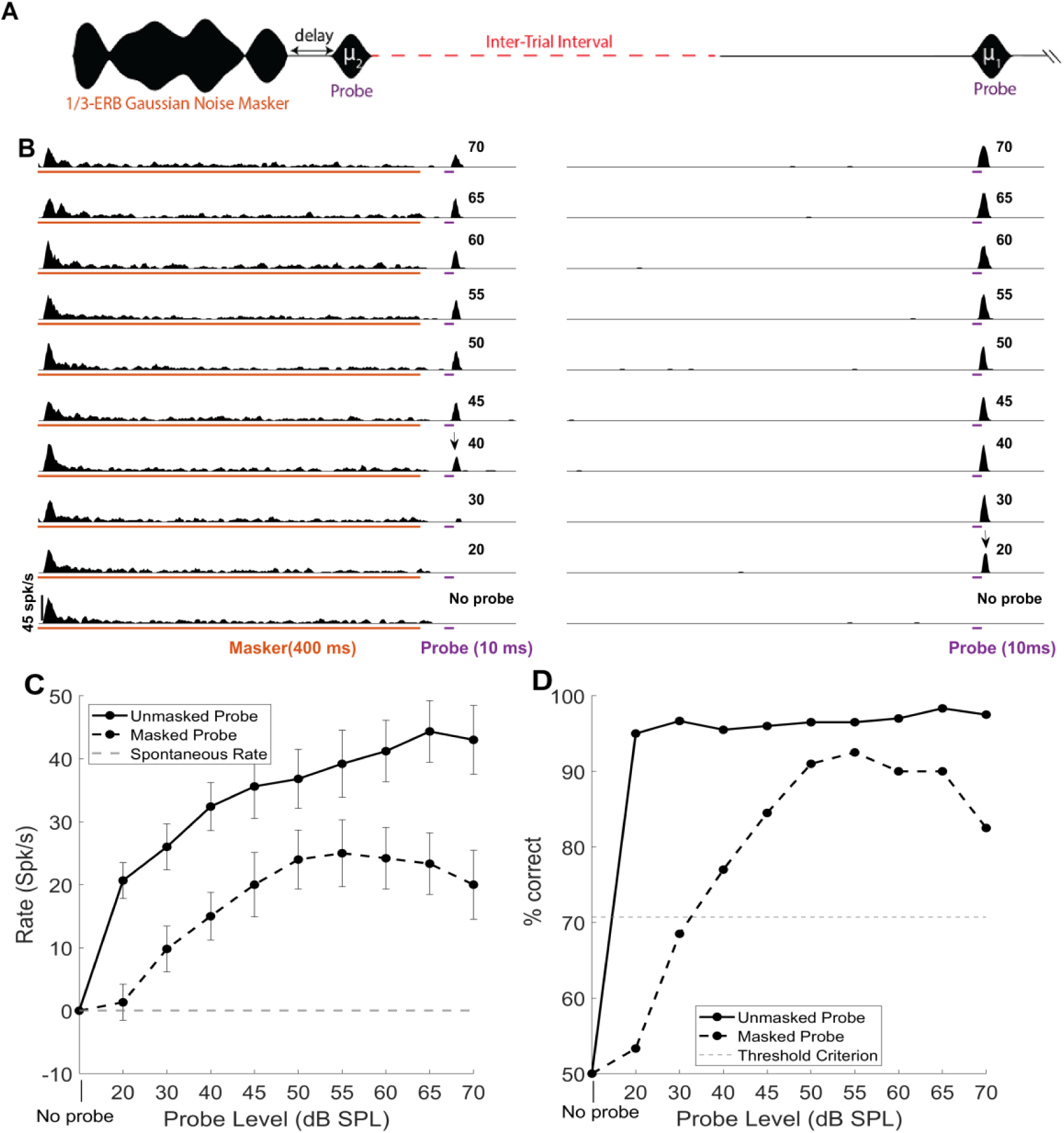
A) Schematic of the stimuli used for threshold estimation. Trials with masked probes and unmasked probes were presented in a pseudorandom manner. ITI (red) was varied across blocks of trials. B) Example PSTHs (1-ms binsize); the lowest probe level at which a neuron responded significantly above the spontaneous rate is indicated by a black arrow for masked and unmasked probes. C) The average and standard deviation of the discharge rate of the neuron in B) for masked (black dashed) and unmasked probes (black solid), with spontaneous activity indicated by the dashed gray line. D) Neurometric function, or percent correct as a function of probe level.

The mean masking levels for single neurons were 16.5 ± 2, 18.5 ± 3, and 21.7 ± 3 dB (mean ± SEM) for ITIs of 0.5, 1, and 1.5 s respectively. The maximum amounts of masking for ITIs of 0.5, 1, and 1.5 s were 45, 55, and 50 dB, respectively, for a 70-dB SPL GN masker with 25-ms delay between masker and probe. The amount of masking did not differ significantly with ITI (Fig. 8).

**Figure 8:**
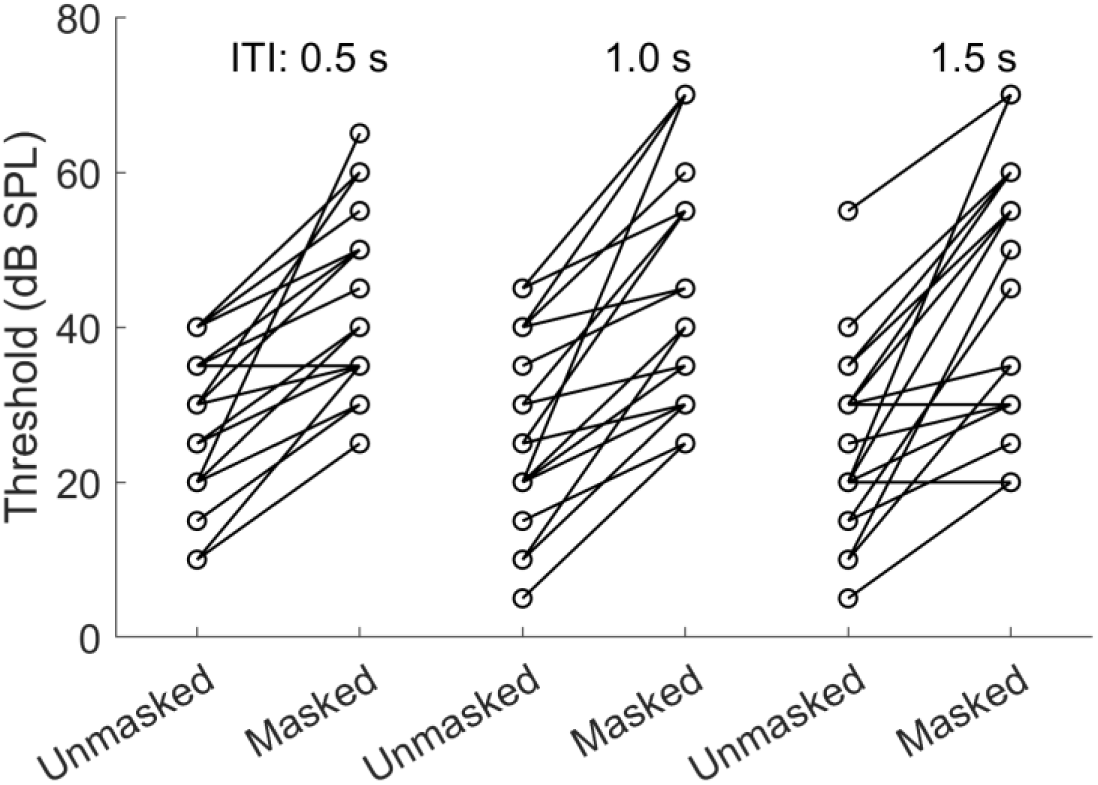
Thresholds for masked and unmasked probes for different ITIs. Each circle represents a neuron’s threshold in the unmasked condition, connected to its threshold in the masked condition at ITIs of 0.5, 1, and 1.5 seconds.

The amount of masking was also estimated for the population of neurons using a maximum-likelihood pattern-decoder analysis of the population response (including n=54, 52, 54 units for ITIs of 0.5, 1 and 1.5 s, respectively) (Day and Delgutte, 2013; Jazayeri and Movshon, 2006). Similar to the results based on the single-unit analysis, the population analysis showed that the amount of masking did not differ significantly with ITI (20+/-1, 20+/-2, 19+/-2 dB +/- sem for ITIs of 0.5, 1, and 1.5 s, respectively).

### E. Testing possible mechanisms for forward masking with GN maskers: Persistence and Medial-olivocochlear efferent gain control

These results provide a basis for testing potential mechanisms for forward masking that have been previously proposed. The psychophysical persistence model of forward masking (Moore et al., 1988; Oxenham, 2001) hypothesizes that masking occurs due to the sustained response to the masker, which overlaps with the response to the probe. To test for physiological evidence of persistence, the discharge rate in the probe-analysis window (a 30-ms window starting at probe onset) for the masker alone condition was compared to the spontaneous rate (calculated during the last 400 ms of the ITI). Responses collected in Experiment 2 were used for this analysis because they included trials with the masker only. Results showed that about 88% of neurons (n=54, 52 and 54 neurons were tested at ITIs of 0.5, 1.0 and 1.5 s, respectively) did not have significant differences in rate between the probe-analysis window for masker-alone trials and spontaneous rate (2-tailed t-test, p > 0.05). A neuron with rate that was significantly different from spontaneous rate using 2-tailed test was then subjected to 1-tailed t-test to determine if spontaneous rate was increased or decreased by the masker. Only 12% of neurons had a significantly increased rate in the probe-analysis window compared to spontaneous activity (1-tailed t-test, p < 0.05). Notably, unlike the auditory nerve, the majority of IC neurons exhibit very low baseline spontaneous rates, with 96% lower than 20 spikes per second. This finding suggests that, similar to the Nelson et al. (2009) results for tone maskers, the majority of IC neurons did not demonstrate masker-response persistence.

Recent computational modeling studies have explored the role of cochlear gain control via the medial-olivocochlear efferent pathway as a potential mechanism that may contribute to forward masking (Brennan et al., 2023; Maxwell et al., 2024). These studies provide insights into psychophysical and physiological aspects related to tone and noise forward maskers, such as suppression of probe responses observed physiologically (Nelson et al. 2009) and robustness to randomly varying masker levels in human listeners (Jesteadt et al. 2005), which are not explained by the persistence model. To determine if efferent control of cochlear gain offers a potential explanation for masking recovery observed in physiological experiments, a computational model with efferents was used to simulate IC responses to GN maskers and probes, similar to those used in Exp. 1.

The response to an unmasked probe (Fig. 9A) and a masked probe (Fig. 9B) for a computational model with efferents, for a delay of 25 ms between the GN masker and probe, shows mean responses of r1 and r2, respectively. The recovery ratio between the masked and unmasked responses (r2/r1) was calculated for different delays, both with and without efferent gain for a CF OF 2000 Hz (Fig. 9C). The masked-probe response of the BE IC model without efferent gain control (dashed blue) was decreased for 0-ms delay between the GN masker and probe relative to the unmasked condition. However, for larger delays, the masked-probe response was comparable to the unmasked-probe response. The response of the model with efferent gain control (dot-dashed red) gradually increased as a function of delay (Fig. 9C). The recovery from masking was quantified by the ratio between the mean of the response to the masked and unmasked probes within a 30-ms window beginning at the probe onset, similar to neural data. The recovery from masking increased gradually with the increase in delay for the model with medial-olivocochlear efferent gain control, with an overall time course and gradual recovery that was more comparable to that of the physiological data for delays less than 500 ms (gray, Fig. 9C). Note that the model response at 0-ms delay is influenced by model refractoriness and rate adaptation, as well as by efferent gain control.

**Figure 9:**
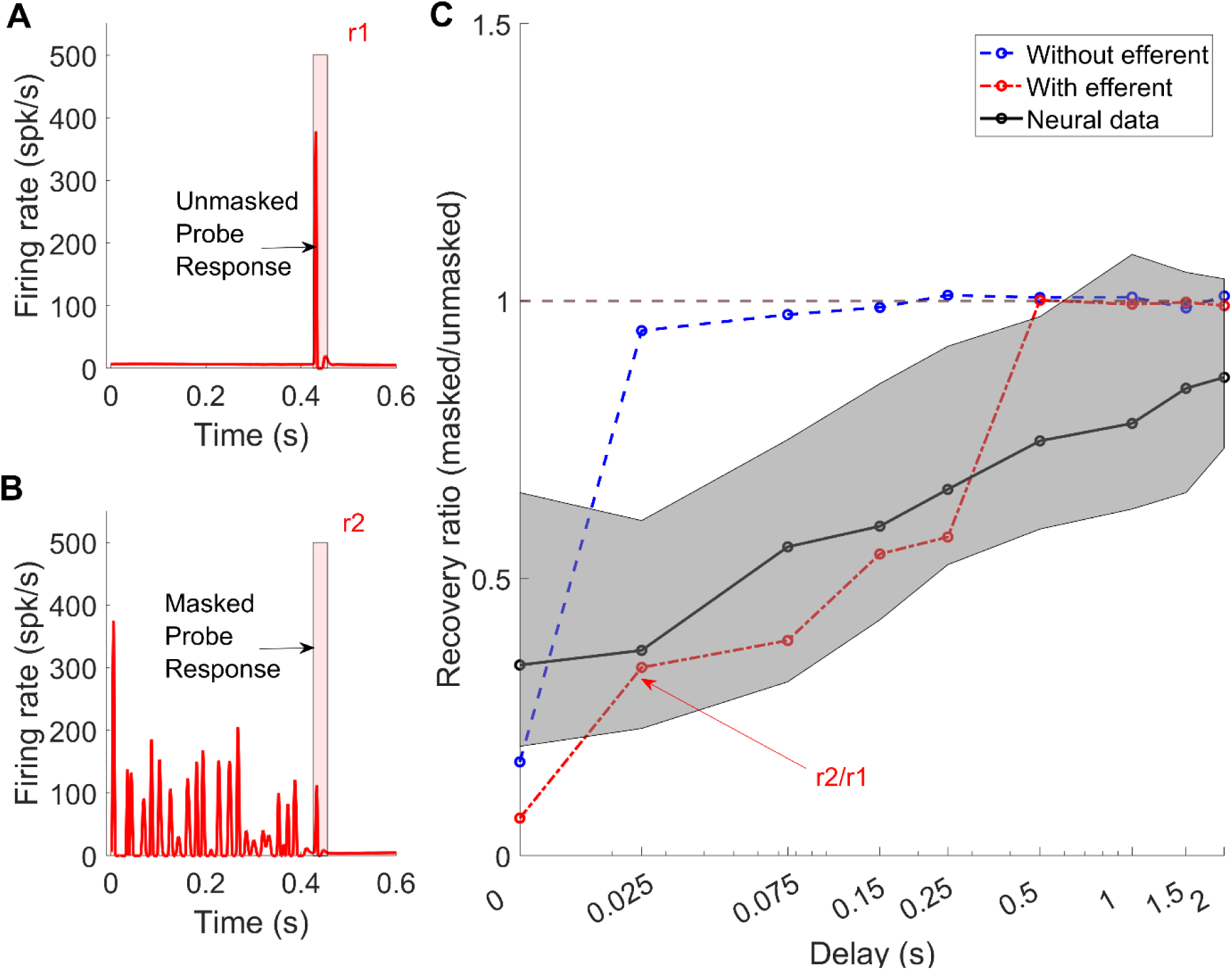
Model responses without (dashed blue) and with (dot-dashed red) efferent control of cochlear gain. A) Responses of AN model with efferents to unmasked and B) masked probes at one delay to illustrate how the mean responses were extracted C) Recovery from masking, the ratio of masked to unmasked probe peak responses, as a function of delay for model responses without (dashed blue) and with (dot-dashed red) efferent gain control. Neural data are shown by the gray shaded area (interquartile range), with medians joined by the black solid line.

## IV. DISCUSSION

Previous physiological studies have explored the impact of tone maskers on probe responses by varying stimulus features such as the delay between the masker and probe or the masker level. The current study explored the recovery of neuronal discharge rate after narrowband-noise forward maskers. In the current study, we observed that a delay greater than 1.5 s was necessary for more than 50% of the neurons to recover after 70-dB SPL GN maskers, a duration significantly longer than the reported recovery times for tone maskers, which typically fall within the range of 100-300 ms (for masker levels 30-60 dB above neural thresholds determined using response maps). More neurons recovered as the delay was increased between the masker and the probe. Neurons with higher CFs tended to have longer recovery times than low-CF neurons, but there was not a significant relationship between spontaneous rate or masker-response type and recovery time.

Furthermore, a small but significant carryover of masking from previous trials, for ITIs less than 1.5 s, was observed by analyzing responses to subsequent unmasked probes. The narrowband GN maskers elevated thresholds by 20-50 dB, and the amount of masking was not significantly affected by ITI.

Physiological studies have examined the relationship between recovery from forward masking and spontaneous activity at different levels of the auditory pathway. Relkin & Doucet (1991) showed that LSR AN fibers tend to exhibit more masking than HSR fibers. Shore (1995) also reported that neurons in the ventral cochlear nucleus with low and medium spontaneous activity had more forward masking than those with high spontaneous activity. Shore reported that the effect is especially evident at high masker levels and short masker-probe intervals, particularly for short (10-ms) probe delays. Nelson et al. (2009) also reported a weak but significant dependence of the amount of masking on spontaneous rate in the IC of awake marmosets for a 10-ms probe delay. In the current IC study, a significant correlation was not observed between the recovery from masking and spontaneous rate, even at a delay of 0 s, where masking was strongest (Fig.5A). The difference between this observation and Nelson et al. (2009) might be attributed to differences in the range of spontaneous activity and species; Nelson et al. (2009) reported spontaneous rates up to 100 spikes/sec in marmoset, whereas the highest spontaneous rate in the rabbit dataset reported here was 40 spikes/sec.

Psychoacoustic studies have explored the frequency dependence of recovery from masking, revealing conflicting results. Some studies (Harris et al., 1951; Luscher and Zwislocki, 1949; Rawnsley and Harris, 1952) indicate longer recovery from masking at high frequencies, while others report longer recovery at low frequencies (Jesteadt et al., 1982). Shore’s (1995) physiological investigation in the ventral cochlear nucleus demonstrated that high-CF neurons were less recovered at short delays, consistent with our results in IC (Fig. 5B). We also found that recovery from masking was independent of whether neurons had responses that increased or decreased over the time course of the masker (Fig. 5C, D). Qualitatively, the relationship of recovery from forward masking and masker-response slope did not vary with RM (Fig. 5C) or MTF type (Fig. 5D).

In the current study, the GN masker level was always fixed at 70 dB SPL and the maximum amount of forward masking was 50 dB (Fig. 8). Psychoacoustic studies have reported forward masking by tone maskers of 15-55 dB (e.g., for a 2-kHz, 100-dB SPL masker, ∼15 dB and ∼55 dB masking was reported for 40- and- and 2-ms probe delays, respectively, Fig. 2 in Plack & Oxenham, 1998; for a 125 Hz, 90-dB SPL masker, up to ∼40 dB of masking was reported for probe delays of 5, 10, 20 ms, Fig. 3 in Jesteadt, 1982). For GN maskers with 1/3-ERBn bandwidths, as used in the current study, a threshold shift of ∼40 dB was observed for a delay of 25 ms and a masker level of 80 dB SPL in young normal hearing listeners (Brennan et al., 2023). Similarly, physiological studies in several species have reported the amount of masking at different locations along the auditory pathway for tone maskers 40 dB above unmasked threshold (auditory cortex, guinea pig, maximum ∼50 dB, Alves-Pinto et al., 2010; medial nucleus of the trapezoid body, rat, maximum ∼54 dB, Gao & Berrebi, 2016; ventral cochlear nucleus, guinea pig, maximum ∼33.5 dB, Ingham et al., 2016; IC, marmoset, maximum ∼50 dB, Nelson et al., 2009; auditory nerve, chinchilla, maximum ∼21 dB forward masking, Relkin & Doucet, 1991.) The disparities in masking levels could be attributed to different recording locations and/or species. Notably, the study by Nelson et al. (2009) was the only previous study to record from the IC of an awake animal (marmoset), while all other studies involved anesthesia.

According to the psychophysical persistence model for forward masking (Moore et al., 1988; Oxenham, 2001), masking arises due to a sustained response to the masker, which overlaps with the response to the probe. However, the physiological study in the IC by Nelson et al. (2009) showed that, for tone maskers, the masker responses were mostly adapted over the course of the masker duration, and there was no significant difference between the spontaneous rate and the average rate during the probe-analysis window in the masker-alone condition. We employed the same metric and found a similar result: forward masking by GN was observed at the level of the IC, but it was not attributable to the persistence of the masker responses for the majority of IC neurons. Note that although both this study and Nelson et al. (2009) observed forward masking at the level of the IC with properties comparable to those observed psychophysically, it would still be interesting to study forward-masking paradigms at higher levels in the auditory pathway in awake animals.

Comparing human forward-masking data using narrowband noise maskers with physiological data is challenging due to the influence of confusion effects on human masked thresholds. Confusion effects occur in psychophysical forward-masking conditions in which listeners struggle to differentiate the signal from the preceding masker (Moore & Glasberg, 1982, 1985; Neff & Jesteadt, 1983; Neff, 1986). These effects are particularly prominent when the masker-signal interval is short, the masker is narrowband noise (50–60 Hz wide), and the signal duration aligns with the average rate of envelope fluctuations in the masker (Moore & Glasberg, 1982, 1985; Neff & Jesteadt, 1983; Neff, 1986). Under these conditions, listeners may find it difficult to perceive the transition between the masker and the signal. As a result, signal thresholds measured in the presence of confusion effects can be artifactually elevated, because the signal is detectable but cannot be reliably distinguished from the masker due to spectral or temporal similarities. In contrast, such issues do not arise in the physiological data presented here, because the observation intervals were explicitly defined relative to the timing of the probe. This key difference complicates direct comparisons between physiological and psychophysical data.

Previous studies (Moore & Glasberg, 1982, 1985; Neff & Jesteadt, 1983) have provided clear evidence of significant confusion effects with narrowband maskers (50–60 Hz wide) centered on the signal frequency. Introducing a contralateral or low-level ipsilateral broadband noise led to threshold reductions of up to 27 dB compared to thresholds with the masker alone. Similarly, contralateral sinusoids also reduced thresholds, with even greater reductions observed under ipsilateral conditions (Moore & Glasberg, 1982). To understand the contribution of confusion effects, future physiological experiments could measure thresholds with and without a temporal cue presented to the ear contralateral to the forward masker and signal.

Alternative mechanisms that may explain forward masking include suppression of the probe response in the IC, arising from the superior para-olivary nucleus in the brainstem (Felix et al., 2015; Gai, 2016; Gao and Berrebi, 2016; Gao et al., 2017; Nelson et al., 2009; Salimi et al., 2017). However, chemical inactivation of the superior para-olivary nucleus does not remove suppression of probe responses (Felix et al., 2015). The role of the efferent system in forward masking has been previously suggested (Jennings, 2021; Krull and Strickland, 2008; Maxwell et al., 2024; Relkin and Turner, 1988). This mechanism was initially discounted because the growth of masking in the AN response, i.e., increased probe threshold with increasing masker level, which was presumed to include the effects of the efferent system, is insufficient to account for psychophysical growth of forward masking.

However, the fact that anesthesia (Guitton et al. 2004) can suppress the efferent system was not taken into account in this argument. In the current study, the recovery from masking as a function of delay was better explained by a subcortical model that included cochlear gain control by the medial-olivocochlear efferent pathway as compared to a model without efferent gain control (Fig. 9B). The model with efferent gain control could account for recovery from masking for delays less than 500 ms, but not for longer delays. This limitation of the model might be due to the fact that it does not include very slow efferent effects (Cooper and Guinan, 2003). The physiological data for the long-term recovery provides a database for developing future models that include longer recoveries, possibly due to slow efferent effects.

This study explored the effects on forward masking of masker-probe delay and ITI for a narrowband GN masker at 70 dB SPL. Other parameters, such as masker duration, probe duration, and masker level remain unexplored; studies of these parameters could provide further insight into differences between forward masking by tones and noise, along with the impact of ITI on forward masking.

## ACKNOWLEDGEMENTS

The work was supported by National Institutes of Health Grant No. R01-DC010813 to LHC. We thank Kristina Abrams and Johanna Fritzinger for their help with the physiological experiments, and Douglas Schwarz for his assistance with the software for neural data collection.

## AUTHORS DECLARATIONS

### Conflict of Interest

The authors have no conflicts to disclose.

### Ethics approval

This study was approved by the University of Rochester Committee on Animal Resources and conducted in accordance with National Institutes of Health guidelines and ASA Ethical Principles.

### Data Availability

The data that support the findings of this study will be publicly available at https://osf.io/n4m5b/.

### End Notes

1. Note that for two preliminary neurons that were included in the analysis for experiment 1, 15 and 25 repetitions were used.
2. Note that for two preliminary neurons that were included in the analysis for experiment 2, only 10 repetitions were used.

## Notes

### Competing Interest Statement

The authors have declared no competing interest.

## References

Alves-Pinto, A., Baudoux, S., Palmer, A. R., & Sumner, C. J. (2010). Forward masking estimated by signal detection theory analysis of neuronal responses in primary auditory cortex. Journal of the Association for Research in Otolaryngology, 11, 477–494.

Brennan, M. A., Svec, A., Farhadi, A., Maxwell, B. N., & Carney, L. H. (2023). Inherent envelope fluctuations in forward masking: Effects of age and hearing loss. The Journal of the Acoustical Society of America, 153(4), 1994–1994.

Brosch, M., & Schreiner, C. E. (1997). Time course of forward masking tuning curves in cat primary auditory cortex. Journal of neurophysiology, 77(2), 923–943.

Carney, L. H., Zilany, M. S., Huang, N. J., Abrams, K. S., & Idrobo, F. (2014). Suboptimal use of neural information in a mammalian auditory system. Journal of Neuroscience, 34(4), 1306–1313.

Carney, L. H., & McDonough, J. M. (2019). Nonlinear auditory models yield new insights into representations of vowels. *Attention, Perception*, & Psychophysics, 81, 1034–1046.

Cooper, N. P., & Guinan Jr, J. J. (2003). Separate mechanical processes underlie fast and slow effects of medial olivocochlear efferent activity. The Journal of physiology, 548(1), 307–312.

Day, M. L., & Delgutte, B. (2013). Decoding sound source location and separation using neural population activity patterns. Journal of Neuroscience, 33(40), 15837–15847.

Egan, J. P. (1975). Signal detection theory and ROC analysis. Academic Press. NewYork,

Elliott, L. L. (1969). Masking of tones before, during, and after brief silent periods in noise. The Journal of the Acoustical Society of America, 45(5), 1277–1279.

Farhadi, A., Jennings, S. G., Strickland, E. A., & Carney, L. H. (2023). Subcortical auditory model including efferent dynamic gain control with inputs from cochlear nucleus and inferior colliculus. The Journal of the Acoustical Society of America, 154(6), 3644–3659.

Fan, L., Henry, K. S., & Carney, L. H. (2021). Responses to diotic tone-in-noise stimuli in the inferior colliculus: stimulus envelope and neural fluctuation cues. Hearing research, 409, 108328.

Fan, L., Henry, K. S., & Carney, L. H. (2022). Responses to dichotic tone-in-noise stimuli in the inferior colliculus. Frontiers in Neuroscience, 16, 997656.

Felix, R.A., Magnusson, A.K. and Berrebi, A.S., 2015. The superior paraolivary nucleus shapes temporal response properties of neurons in the inferior colliculus. Brain Structure and Function, 220, pp.2639–2652.

Fishman, Y. I., Arezzo, J. C., & Steinschneider, M. (2004). Auditory stream segregation in monkey auditory cortex: effects of frequency separation, presentation rate, and tone duration. The Journal of the Acoustical Society of America, 116(3), 1656–1670.

Fraisse, P. (1984). Perception and estimation of time. Annual review of psychology, 35(1), 1–37].

Gai, Y. (2016). ON and OFF inhibition as mechanisms for forward masking in the inferior colliculus: a modeling study. Journal of Neurophysiology, 115(5), 2485–2500.

Gao, F., & Berrebi, A. S. (2016). Forward masking in the medial nucleus of the trapezoid body of the rat. Brain Structure and Function, 221, 2303–2317.

Gao, F., Kadner, A., Felix, R. A., Chen, L., & Berrebi, A. S. (2017). Forward masking in the superior paraolivary nucleus of the rat. Brain Structure and Function, 222, 365–379.

Glasberg, B. R., & Moore, B. C. (1990). Derivation of auditory filter shapes from notched-noise data. Hearing research, 47(1-2), 103–138.

Guitton, M.J., Avan, P., Puel, J.L. and Bonfils, P., 2004. Medial olivocochlear efferent activity in awake guinea pigs. Neuroreport, 15(9), pp.1379–1382.

Guest, D. R., & Carney, L. H. (2024). A fast and accurate approximation of power-law adaptation for auditory computational models. The Journal of the Acoustical Society of America, 156(6), 3954–3957.

Harris, J. D., Rawnsley, A. I., & Kelsey, P. (1951). Studies in short-duration auditory fatigue: I. Frequency differences as a function of intensity. Journal of Experimental Psychology, 42(6), 430.

Hirsh, I. J. (1959). Auditory perception of temporal order. The Journal of the Acoustical Society of America, 31(6), 759–767.

Ingham, N. J., Itatani, N., Bleeck, S., & Winter, I. M. (2016). Enhancement of forward suppression begins in the ventral cochlear nucleus. brain research, 1639, 13–27.

Jazayeri, M., & Movshon, J. A. (2006). Optimal representation of sensory information by neural populations. Nature neuroscience, 9(5), 690–696.

Jesteadt, W., Bacon, S. P., & Lehman, J. R. (1982). Forward masking as a function of frequency, masker level, and signal delay. The Journal of the Acoustical Society of America, 71(4), 950–962.

Jennings, S. G. (2021). The role of the medial olivocochlear reflex in psychophysical masking and intensity resolution in humans: A review. Journal of Neurophysiology, 125(6), 2279–2308.

Kaltenbach, J. A., Meleca, R. J., Falzarano, P. R., Myers, S. F., & Simpson, T. H. (1993). Forward masking properties of neurons in the dorsal cochlear nucleus: possible role in the process of echo suppression. Hearing research, 67(1-2), 35–44.

Kidd Jr, G., & Feth, L. L. (1982). Effects of masker duration in pure-tone forward masking. The journal of the Acoustical Society of America, 72(5), 1384–1386.

Kim, D. O., Carney, L., & Kuwada, S. (2020). Amplitude modulation transfer functions reveal opposing populations within both the inferior colliculus and medial geniculate body. Journal of Neurophysiology, 124(4), 1198–1215.

Krull, V., & Strickland, E. A. (2008). The effect of a precursor on growth of forward masking. The Journal of the Acoustical Society of America, 123(6), 4352–4357.

Lisker, L., & Abramson, A. S. (1964). A cross-language study of voicing in initial stops: Acoustical measurements. Word, 20(3), 384–422.

Lüscher, E., & Zwislocki, J. (1949). Adaptation of the ear to sound stimuli. The Journal of the Acoustical Society of America, 21(2), 135–139.

Maxwell, B.N., Farhadi, A., Brennan, M.H., Svec, A. and Carney, L.H. (2024, in press). Auditory Forward Masking Explained by a Subcortical Model with Efferent Control of Cochlear Gain. eneuro.

Mitchell, P. W., Henry, K. S., & Carney, L. H. (2023). Sensitivity to direction and velocity of fast frequency chirps in the inferior colliculus of awake rabbit. Hearing Research, 440, 108915.

Moore, B. C., & Glasberg, B. R. (1982). Contralateral and ipsilateral cueing in forward masking. The Journal of the Acoustical Society of America, 71(4), 942–945.

Moore, B. C., & Glasberg, B. R. (1983). Growth of forward masking for sinusoidal and noise maskers as a function of signal delay; implications for suppression in noise. The Journal of the Acoustical Society of America, 73(4), 1249–1259.

Moore, B. C., & Glasberg, B. R. (1985). The danger of using narrow-band noise maskers to measure ‘‘suppression’’. The Journal of the Acoustical Society of America, 77(6), 2137–2141.

Moore, B. C., Glasberg, B. R., Plack, C. J., & Biswas, A. K. (1988). The shape of the ear’s temporal window. The Journal of the Acoustical Society of America, 83(3), 1102–1116.

Neff, D. L., & Jesteadt, W. (1983). Additivity of forward masking. The Journal of the Acoustical Society of America, 74(6), 1695–1701.

Neff, D. L. (1986). Confusion effects with sinusoidal and narrow-band noise forward maskers. The Journal of the Acoustical Society of America, 79(5), 1519–1529.

Nelson, P. C., Smith, Z. M., & Young, E. D. (2009). Wide-dynamic-range forward suppression in marmoset inferior colliculus neurons is generated centrally and accounts for perceptual masking. Journal of Neuroscience, 29(8), 2553–2562.

Nelson, P. C., & Carney, L. H. (2004). A phenomenological model of peripheral and central neural responses to amplitude-modulated tones. The Journal of the Acoustical Society of America, 116(4), 2173–2186.

Oxenham, A. J., & Plack, C. J. (2000). Effects of masker frequency and duration in forward masking: further evidence for the influence of peripheral nonlinearity. Hearing research, 150(1-2), 258–266.

Oxenham, A. J. (2001). Forward masking: Adaptation or integration? The Journal of the Acoustical Society of America, 109(2), 732–741.

Plack, C. J., & Oxenham, A. J. (1998). Basilar-membrane nonlinearity and the growth of forward masking. The Journal of the Acoustical Society of America, 103(3), 1598–1608.

Plomp, R. (1964). Rate of decay of auditory sensation. The Journal of the Acoustical Society of America, 36(2), 277–282.

Ramachandran, R., Davis, K. A., & May, B. J. (1999). Single-unit responses in the inferior colliculus of decerebrate cats I. Classification based on frequency response maps. Journal of neurophysiology, 82(1), 152–163.

Rawnsley, A. I., & Harris, J. D. (1952). Studies in short-duration auditory fatigue: II. Recovery time. Journal of Experimental Psychology, 43(2), 138.

Relkin, E. M., & Turner, C. W. (1988). A reexamination of forward masking in the auditory nerve. The Journal of the Acoustical Society of America, 84(2), 584–591.

Relkin, E. M., & Doucet, J. R. (1991). Recovery from prior stimulation. I: Relationship to spontaneous firing rates of primary auditory neurons. Hearing research, 55(2), 215–222).

Salimi, N., Zilany, M. S., & Carney, L. H. (2017). Modeling responses in the superior paraolivary nucleus: implications for forward masking in the inferior colliculus. Journal of the Association for Research in Otolaryngology, 18, 441–456.

Schwarz, D. M., Zilany, M. S., Skevington, M., Huang, N. J., Flynn, B. C., & Carney, L. H. (2012). Semi-supervised spike sorting using pattern matching and a scaled Mahalanobis distance metric. Journal of neuroscience methods, 206(2), 120–131.

Shore, S. E. (1995). Recovery of forward-masked responses in ventral cochlear nucleus neurons. Hearing research, 82(1), 31–43.

Stilp, C. (2020). Acoustic context effects in speech perception. Wiley Interdisciplinary Reviews: Cognitive Science, 11(1), e1517.

Turner, C. W., Relkin, E. M., & Doucet, J. (1994). Psychophysical and physiological forward masking studies: Probe duration and rise-time effects. The Journal of the Acoustical Society of America, 96(2), 795–800.

Weber, D. L., & Moore, B. C. (1981). Forward masking by sinusoidal and noise maskers. The Journal of the Acoustical Society of America, 69(5), 1402–1409.

Whitehead, M. L., Lonsbury-Martin, B. L., & Martin, G. K. (1992). Evidence for two discrete sources of 2 f 1− f 2 distortion-product otoacoustic emission in rabbit. II: Differential physiological vulnerability. The Journal of the Acoustical Society of America, 92(5), 2662–2682.

Von Steinbüchel, N., & Pöppel, E. (1991). Temporal order threshold and language perception. Frontiers in knowledge-based computing. Narosa Publishing House, 81–90.

Zilany, M. S., Bruce, I. C., & Carney, L. H. (2014). Updated parameters and expanded simulation options for a model of the auditory periphery. The Journal of the Acoustical Society of America, 135(1), 283–286.

